# Movement responses to lethal risk: an integrative analysis of proactive and reactive antipredator behaviours in a large herbivore

**DOI:** 10.1101/2024.08.03.606415

**Authors:** Charlotte Vanderlocht, Benjamin Robira, Andrea Corradini, Simone Dal Farra, Federico Ossi, Davide Righetti, Heidi C. Hauffe, Luca Pedrotti, Francesca Cagnacci

## Abstract

Prey species can display antipredation movement behaviours to reduce predation risk including proactive responses to chronic or predictable risk, and reactive responses to acute or unpredictable risk. Thus, at any given time, prey movement choice may reflect the trade-off between proaction and reaction. In previous studies, proaction and reaction have generally been considered separately, which neglects their simultaneous influence on animal decisions. In this study, we analysed how proaction and reaction interact to shape the movements of GPS-collared red deer (*Cervus elaphus*), in response to human hunting of conspecifics. Our results show that red deer proactively selected canopy cover where and when risk was predictably high. However, when they were unable to avoid risk, canopy cover was no longer selected, but only modulated a reactive response along a freeze-to-escape continuum. This reaction was even more evident when the environment was unfamiliar, underlining the importance of memory in such reaction patterns. Therefore, to our knowledge, for the first time, we describe how proaction and reaction fuse in an *antipredation sequence* of interconnected movement decisions in a large herbivore, and we lay the foundations for further investigations into the evolutionary origins of similarities and differences between proactive and reactive behaviours.

## 1. Introduction

Avoiding predation while ensuring sufficient resource acquisition is a constant challenge for wild prey species. As prey seek feeding opportunities, they often expose themselves to higher predation risk (Sih 1980) as predators, in turn, also seek high fitness gains by hunting when and where encounter and capture probabilities are high (Mitchell & Lima 2002, Iorio-Merlo et al. 2022). Antipredator behaviours may reduce predation risk, but these are offset by other fitness-related costs (Ajie et al. 2007) such as reduced reproduction (Creel et al. 2007, Zanette et al. 2011), slower growth (Pangle et al. 2007) and/or decreased survival (Griffin et al. 2011). Ultimately, prey individuals must continuously make cost-benefit assessments (Lima & Dill 1990, Brown et al. 1999), trading-off between predation avoidance and resource acquisition.

Empirical measurements of this ‘food-risk trade-off’ in wild populations can be challenging (Jessup & Bohannan 2008); therefore, they are often estimated using unidimensional indices (Kneitel & Chase 2003) such as vigilance levels (Fortin et al. 2004) or ‘giving-up densities’ of resources (Brown 1988). A more comprehensive approach to investigating spatial avoidance despite resource availability according to spatially varying risk or perceived risk has been developed (the ‘landscape of fear’; Laundré et al. 2001; see Gaynor et al. 2019 for review), but these studies often overlook the temporal variability of risk (Palmer et al. 2022). Experimental setups of community compositions, such as control-treatment mesocosm experiments (Sih 1992, Schmitz et al. 1997), have been used to investigate behavioural implications of predator presence on prey species over time, but in spatially small and simplified risk contexts (i.e. presence/absence of predator). However, how the broader spatiotemporal context and fine-scale perception of risk simultaneously combine to exert individual behavioural decisions in a food-risk trade-off is less frequently addressed.

For large vertebrates, this gap in knowledge may be bridged by movement analysis (Nathan et al. 2008), as movement inherently tracks the links between fine-scale behaviours, space use, short and long term changes in external conditions, and thus, context-specific decision-making. Furthermore, movement paths are spatiotemporally scalable, as each movement phase extracted from individual trajectories reflects the behavioural decision of the animal in a specific environment, in a specific internal state, with a specific history of past decisions, and at a specific temporal scale (Nathan et al. 2008, Benhamou 2014). This spatiotemporal scalability of movement paths can be especially relevant in the analysis of antipredator behaviour, as predation can also be segmented into various steps at different spatiotemporal scales (the ‘predation sequence’: encounter - detection - identification - attack - subjugation - consumption; Endler 1991). Antipredator movement behaviours can target various steps of the predation sequence (Endler 1991, Patin 2020): *proactive* antipredator behaviours generally aim at reducing encounter and/or detection probabilities based on cumulative former experience of risk, which may be perceived as chronic in the landscape (Laundré et al. 2010), predictably variable in time (Lima & Bednekoff 1999, Tilmann 2009) and/or predictably associated with environmental contexts (e.g. habitat type: Creel et al. 2014; habitat feature: Kuijper et al. 2015). On the other hand, *reactive* antipredator behaviours generally aim at reducing attack and/or kill probabilities based on direct, sudden cues of risk which are acute or unpredictable to the prey (Creel et al. 2018, Palmer et al. 2022; e.g. Creel et al. 2005, Chassagneux et al. 2020). Moreover, the display of these antipredator behaviours in prey species requires sensory and cognitive abilities (Dall et al. 2005). Indeed, reactive behaviour involves the perception of risk cues while proactive behaviour additionally requires associations between risk and context (e.g. risky time of day, risky habitat feature, risky space), the ‘knowledge’ of which can come from past experience (Fraker 2009) and memory thereof (Bracis et al. 2018), or from genetic priors (Chamaillé-Jammes et al. 2014, Hébert et al. 2019). Furthermore, familiarity with the surrounding area (Clarke et al. 1993, Forrester et al. 2015) can modulate habitat selection (Wolf et al. 2009) and behaviour choice (Chassagneux et al. 2020).

The success of proactive and reactive antipredator behaviours depends on the hunting strategy of the predator (Palmer & Packer 2021). For example, coursing predators tend to increase preys’ perception of risk in open areas, while ambush predators tend to increase the perception of risk in covered habitat (Chen et al. 2021). Human hunters, while exhibiting a large diversity in hunting modes, select areas with good visibility to improve hunting success (Gaynor et al. 2024). Human disturbance is also a form of risk (Frid & Dill 2002, Ciuti et al. 2012), and ungulates in anthropogenic landscapes are known to switch between forested areas during the day, and open areas at night (Salvatori et al. 2022). Therefore, in ungulates that coevolved with coursing predators such as wolves (*Canis lupus*), and/or with human hunters and disturbance, the food-risk trade-off is often translated into a compromise between grazing in open areas, where grass biomass (Ratajczak et al. 2012, Randle et al. 2018) and palatability (Jefferies et al. 1994) may be higher, and selecting canopy cover to evade predators (Mysterud & Østbye 1999). Canopy cover can be proactively selected, in response to predictable or chronic risk, but the role of canopy cover in antipredation goes beyond simple selection or avoidance; for example, perception of risk in open habitat can be exacerbated by the distance to refuge cover (Stankowich 2008), or by the relative positions between the predator, the prey and the refuge (Kramer & Bonenfant 1997). When exposed to direct and acute lethal risk, the classical escape *vs* freeze decisional dilemma (Eilam 2005) that ungulates face (Stankowich 2008, Chassagneux et al. 2019) can also be modulated by cover (Bonnot et al. 2017). Ungulates therefore use canopy cover as a proactive *and* as a reactive antipredator behaviour.

Proaction and reaction coexist within the spectrum of behaviours available to prey species, but little is known about how the costs and benefits of these often presumed independent antipredator behaviours are associated, and how they impact an individual’s decision-making and movement path, in relation to canopy cover and lethal risk. To fill this gap, we used high-resolution hunting data combined with hourly movement data of GPS-collared red deer (*Cervus elaphus*) from the same area. Specifically, we investigated the role of canopy cover in a complex anthropogenic Alpine environment in shaping these antipredator responses, providing a deeper understanding of antipredator movement behaviours along the predation sequence and at multiple spatiotemporal scales (Fig.1).

**Figure 1.**
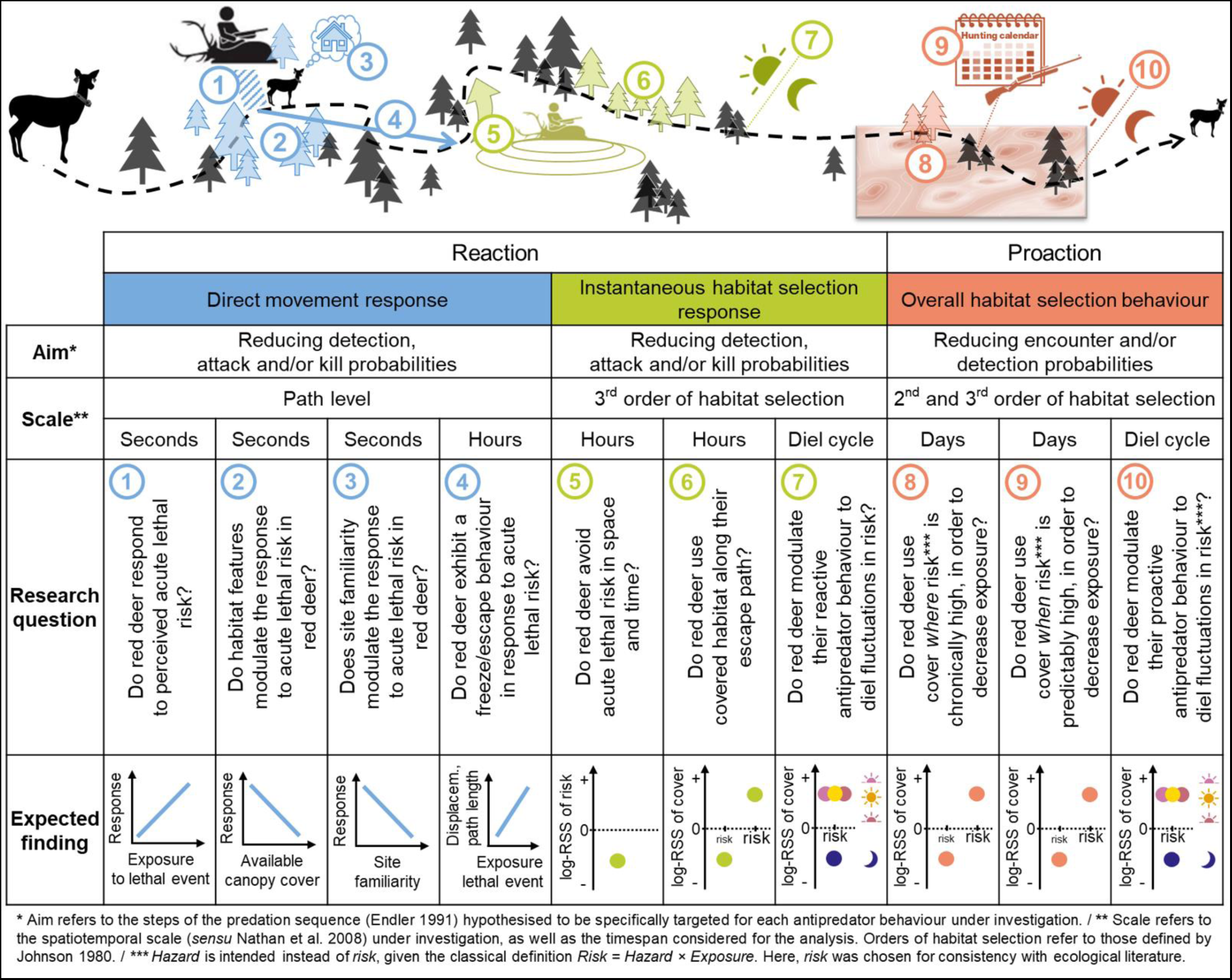
Schematic representation of predicted associations between predation risk and prey movement along the predation sequence, including our research questions and expectations.

We considered three main research questions:

- **How do red deer respond to acute lethal risk through movement, given exposure to the risk, surrounding habitat features and site familiarity? (Q.1-4)** Given the results of Chassagneux et al. (2020) showing that ‘freezing’ in red deer was always followed by delayed flight, we expect that red deer ‘escape’ after closely encountering and perceiving lethal risk. We thus expect that red deer perceive hunting events in their direct surroundings, and that they respond to these events by modifying movement metrics in the hours following the encounter, with a decreasing response magnitude with distance to the hunting location, (Q.1) as (signals of) risk dilute(s) in space; with available surrounding canopy cover (Q.2), as refuge habitat may reduce (perceived) risk levels; and with site familiarity (Q.3), as knowledge of the surrounding areas may make refuge seeking more efficient. Specifically, we expect that red deer exhibit an ‘escape’ response in the 24 h following exposure to direct lethal risk (Q.4), characterised by increased displacement and increased cumulated path length.
- **What is the instantaneous habitat selection response of red deer to acute lethal risk, given the diel period? (Q.5-7)** If red deer not only adjust the way they move as an immediate response to hunting risk, but also where they move, we expect red deer to (Q.5) avoid acute lethal risk. When exposed to high acute risk, we expect they will (Q.6) select for forested habitat given the role of canopy cover as a refuge from human hunters and disturbance. We expect these reactive antipredator behaviours to be (Q.7) strongest during the hours of the day in which hunting is allowed (i.e. dawn, day, dusk).
- **What is the overall habitat selection response of red deer in a landscape of predictable spatial and temporal risk? (Q.8-10)** If red deer adjust their movement behaviours to chronic or predictable risk in the area (i.e. proactive behaviour), we expect that (Q.8) red deer use cover where chronic spatial risk is high in order to decrease exposure, and that (Q.9) they increase their selection for cover as hunting pressure increases throughout the year (i.e. during the hunting season). Finally, we also expect that (Q.10) proactive antipredator behaviour in red deer is strongest during the day when hunting is allowed and human disturbance is prevalent. Given the crepuscular nature of red deer (Ensing et al. 2014), and the periodic feeding needs of ruminants (Hofmann 1989), we also expect that (Q.10) red deer might take more risks at dawn and at dusk despite hunter activity.

## 2. Materials and methods

### 2.1. Study area and red deer hunting regime

The study took place in the hunting reserves of Lauregno and Proves, in the Non Valley (Central-eastern Alps, Autonomous Province of Bolzano, Italy), which cover a total surface of 32.7 km^2^ (Fig.S1), characterised by rugged terrain (857-2613 m a.s.l.). Lower altitudes are dominated by conifer forests (*Abies spp.* and *Picea spp.*), as well as irrigated hay meadows, while higher elevations are covered successively by alpine shrublands and grasslands. The grey wolf (*Canis lupus*) has been recolonising the area since 2017 (Groff et al. 2018). The population of red deer in this study area has been estimated as approximately 278 individuals (average number of individuals counted between 2022 and 2024 in the Lauregno and Proves preserves; source: Autonomous Province of Bolzano), and is selectively hunted (hunting quotas attain *circa* 45 heads per year in Lauregno and *circa* 20 heads per year in Proves) following the Italian law and regulations of the Province of Bolzano (https://provincia.bz.it/agricoltura-foreste/fauna-caccia-pesca/default.asp): that is, selective hunting is open from May 1st to December 15th, with demographic targets varying throughout the year. Hunting is banned at night (one hour after sunset to one hour before sunrise). Ungulate hunters use firearms, and adopt a sit-and-wait approach, at ground level or using an elevated hunting blind.

### 2.2. Red deer captures

We captured 15 adult female red deer during the winters of 2020-2023, reaching a total of 23 animal-years between 11/01/2021 and 20/11/2023, each of which were eartagged and collared with a GPS tracker (Vectronics Aerospace GmbH©, Vertex Plus), set to record one fix/hour, for *circa* 1.5 y after which the collars were programmed to drop off. During captures, deer were immobilised through telenarcosis (‘Vienna mix’, i.e. Xylazine, Zoletil, and Antisedan). All procedures were performed with ethical permission (permit approval by ISPRA-Wildbeobachtungsstelle) and led by a licensed veterinarian following Italian regulations. The administrative authorisation was granted by Decree No. 24664/2020 of the competent Provincial Council.

### 2.3. Data pre-processing

We carried out data processing and analyses in R software (v.4.2.2; R Core Team, 2022; see Data S2 for the list of used R packages).

#### 2.3.1. Trajectory data

We processed the GPS data by removing locations with low accuracy (dilution of precision > 10), and added 10 s to time duplicates and 0.00001 degrees latitude and longitude to location duplicates. In order to exclude any capture effect from the analysis, we also removed all locations in the 10 days following capture (Bergvall et al. 2021). In the resulting animal tracks, 83% of all fixes were separated from the previous and subsequent fix by 1 h (± 10 min). We did not interpolate missing locations, to only focus on ‘true’ locations. After data pre-processing, a total of 137,368 GPS locations were used for subsequent analyses (Fig.S1).

#### 2.3.2. Hunting data

All legal hunting events were recorded in the database of the South Tyrol Alto Adige Hunters’ Association (www.stat.jagdverband.it), with the name of the hunter, species, location (estimated precision of *circa* 15 min) and kill time (estimated precision of *circa* 20 min). In order to account for former experience of risk, as well as hunting events concurrent to red deer data, we extracted the hunting records for all red deer bagged on the territory since 2016 (i.e. the earliest available date; extraction on the 05/02/2024 with *qdapRegex* R package), obtaining a total of 470 reported hunting events (annual mean of 58.8 ± 3.2 events). We excluded incomplete records (e.g. missing timestamp, missing location), reaching a total of 407 red deer hunting events (Fig.S1). These were used to calculate lethal risk variables (see 2.4.2).

### 2.4. Model variables

In order to investigate the role of canopy cover in antipredator movement behaviours of red deer at various spatiotemporal scales (Fig.1), we extracted ecological variables from spatial layers as described below.

#### 2.4.1. Environmental variables

We described the environment of red deer with four spatial environmental layers known to affect red deer movement behaviour in Alpine environments, in raster format at a resolution of 10 m. We derived altitude and terrain ruggedness from the digital elevation model of the National Institute of Geophysics and Volcanology (INGV; Data S3) for topographical description of the area. We generated a canopy cover density map, calculated as the surface density of trees exceeding 2 m height in LiDAR’s Canopy Height Model (Data S3) to describe refuge distribution. As a proxy for quality forage availability, we also estimated daily food distribution maps by combining remote sensing data and field measurements obtained from systematic vegetation surveys (Data S3).

#### 2.4.2. Lethal risk variables

Based on the locations and timestamps of all hunting events in the study area, we derived various components of lethal risk experienced by the collared red deer. First, we quantified *acute* spatiotemporal risk, to investigate reactive responses as a function of spatial and temporal distance to lethal events occurring throughout the monitoring period of red deer (11/01/2021-20/11/2023), considering the closest distance to the kills within one hour prior to the hunting event (to study the direct movement response; Q.1-4), and considering hourly spatially-explicit risk maps (to study the instantaneous habitat selection response; Q.5-7). Second, we quantified *chronic* spatial and *predictable* temporal risk, to investigate proactive movement behaviour as a function of canopy cover and risk itself (overall habitat selection; Q.8-10), by considering the spatial and temporal kernels estimated on all lethal events since 2016.

First, for every hunting event occurring throughout the monitoring period, we first measured the distance between the location of the hunting event and each GPS-collared red deer at the time of hunting (or at the closest fix before the hunting event within 1 h). Second, we created a series of hourly risk maps at a resolution of 10 m. We estimated the weighted kernel density considering hunting events that occurred within the previous 24 h (smoothing parameter of 2 km; *spatialEco* R package; the threshold of 2 km was identified from the predictions of the reaction to lethal risk model, see 3.1; see also Proffitt et al. 2009, Creel et al. 2005). The used weights were a function of time and decreased linearly from 1 to 0 throughout the 24 h following the hunting event, in order to simulate the decay of perceived acute risk. Each hourly risk map was then weighted overall by the highest weight occurring in that map, as a way of distinguishing high and low absolute risk values between hourly maps. Third, to create the chronic spatial risk map, we generated a map of cumulated spatial risk at a resolution of 10 m: we estimated the kernel utilisation distribution (*ad hoc* estimation of smoothing parameter; *adehabitatHR* R package) of all locations of recorded hunting events between January 2016 and November 2023. To estimate the predictable temporal risk, we fitted a yearly circular kernel density (smoothing parameter of 0.1; *circular* R package) to the circular Julian dates of all recorded hunting events between January 2016 and November 2023.

All risk variables were constrained between 0 and 1. We extracted the risk for each location as the density value of the cell containing this location (acute or chronic risk), associated with its timestamp (temporal chronic risk).

#### 2.4.3. Diel periods

We defined four diel periods in the 24 h of each day relative to the timing of sunrise and sunset (extracted with the *suncalc* R package): dawn and dusk were considered as the two-hour periods centred around sunrise and sunset, respectively, and day and night in the remaining hours from dawn to dusk, and from dusk to dawn, respectively. Hunting events occurred during ‘dawn’ (12% of all hunting events), ‘day’ (26%) and ‘dusk’ (62%); but not at ‘night’ (Fig.S4).

### 2.5. Movement analyses

#### 2.5.1. **Reaction to lethal risk**: direct movement response

To characterise red deer movement following a lethal event, we measured red deer displacement (straight line distance between the first and last location) and cumulated path length over the 24 h following the lethal event. We excluded travel sequences with missing hourly fixes in such time intervals. We then investigated how movement metrics (displacement and cumulated path length) varied as a function of risk, using a generalised linear mixed model with a negative binomial error distribution and a log family (*glmmTMB* R package, *nbinom2* as family). Specifically, we modelled these movement metrics as a function of the distance to the lethal event at the time of hunting (Q.1), interacting with the canopy cover density surrounding the animal (buffer of 500 m) at the time of hunting (Q.2), and site familiarity (as the movement-based utilisation distribution value associated with the location at time of the lethal event; Data S5) (Q.3). We also controlled for the local terrain ruggedness (Data S3) at the time and location of hunting, as well as individuals and years as random effects on the intercept. Continuous explanatory variables were scaled and centred (distance to lethal, terrain ruggedness), whereas proportional explanatory variables (canopy cover density, site familiarity) were arcsine-square-root transformed. We analysed the effects of risk and environmental variables on displacement and cumulated path length in two separate models (Q.4), and we verified the models’ assumptions, as well as the fit quality by statistically assessing the prevalence of outliers (Tables S6.2 and S7.2; *DHARMa* R package), and visually checking residual distributions (Fig.S6.3, Fig.S7.3; *DHARMa* R package). We did not note any collinearity issue (maximum variance inflation factor < 2; Tables S6.4 and S7.4; *performance* R package).

#### 2.5.2. Habitat selection

We fitted integrated step selection functions (Avgar et al. 2016) on individual movement tracks to estimate the relative probability of selecting covered habitat at various levels of lethal risk, as well as the relative probability of selecting spatial risk itself, both reactively (Q.5-7) and proactively (Q.8-10). To do so, we generated 50 potential steps (‘available’) for each observed (‘used’). These available steps were obtained by sampling their distances in a Gamma distribution and the turning angles in a Von Mises distribution, parameterised based on observed steps. We then fitted conditional logistic regressions (*amt* R package) on all individual movement choices together, considering risk (see 2.4.2) and environmental variables (see 2.4.1; with values constrained between 0 and 1) as predictors, controlling for movement variables (cosine of the turning angle and logarithm of the length of the step), and we used the individual-step ID as stratum. Due to unavailability of risk information outside of the hunting preserves, we removed ‘used’ steps with missing risk values, or individual-step ID with more than 10 missing ‘available’ risk values. The quantitative effect of variables significantly affecting movement choices were estimated with logarithmic relative selection strengths (log-RSS; Avgar et al. 2017), namely by computing the logarithm of the ratio between the model predictions when all variables were set to 0.5 (exception for Fig.3B, see caption), while the variable of interest was set to 1 or 0, and the step length and turning angle to their mean values.

##### 2.5.2.1. **Reaction to lethal risk:** instantaneous habitat selection response

To model the reactive behaviour of red deer to acute lethal risk, we considered the probability of a location to be selected as a function of the acute spatiotemporal risk (Q.5), the canopy cover density (Q.6) and the diel period (Q.7).

##### 2.5.2.2. **Proaction to lethal risk:** overall habitat selection behaviour

To model the proactive behaviour of red deer to chronic lethal risk, we estimated the probability of a location to be selected as a function of the chronic spatial risk (Q.8), the predictable temporal risk (Q.9), the diel period (Q.10) and the canopy cover density.

For both habitat selection models (reaction and proaction), we ran alternative models including some or all of the above-mentioned variables of interest: from simple additive terms to complex three-way interactions (Tables S8 and S9). We also included control variables to all tested models, namely food availability, altitude and terrain ruggedness as environmental additive fixed terms, and step length in interaction with turning angle as well as step length in interaction with diel period as movement-dependent fixed terms (Avgar et al. 2016). Among the alternative models (Tables S8 and S9), the most parsimonious was selected (Burnham et al. 2011), minimising the Akaike Information Criterion (AIC).

## 3. Results

### 3.1. Reaction to lethal risk: direct movement response

Our models for displacement (displ) and path length (pathl) explained 25% and 27% of variability observed in our data, respectively. Distance to the lethal event in interaction with surrounding canopy cover density strongly and significantly modulated red deer displacement (χ²_1_ = 11.09, p_LRT_ = 0.001) and path length (χ²_1_ = 4.05, p_LRT_ = 0.044) response in the following 24 h. Specifically, when red deer were mainly surrounded by open areas, displacement decreased as distance to the lethal event increased (Fig.2B – dotted red line). In contrast, when red deer were surrounded by canopy cover, displacement increased as distance to the lethal event increased (Fig.2A – solid green line). These opposing behaviours intersected at about 2 km from the lethal event, and at this distance red deer showed the same displacement regardless of the canopy cover. Similar trends were observed for path lengths, with a converging response intensity regardless of surrounding cover only when red deer were very close to the lethal event (< 100 m) (Fig.2C). Displacement and path length were significantly lower when red deer were familiar with their surroundings (Fig.2B,D; *β*_displ_ = −0.370, SE_displ_ = 0.052, *z*_displ_ = −7.09, *p*_displ_ < 0.001; *β*_pathl_ = −0.122, SE_pathl_ = 0.025, *z*_pathl_ = −4.95, *p*_pathl_ < 0.001). Local terrain ruggedness did not affect the direct movement response of the deer (χ²_1,displ_ = 0.25, p_LRT,displ_ = 0.618; χ²_1,pathl_ = 0.34, p_LRT-pathl_ = 0.600).

**Figure 2.**
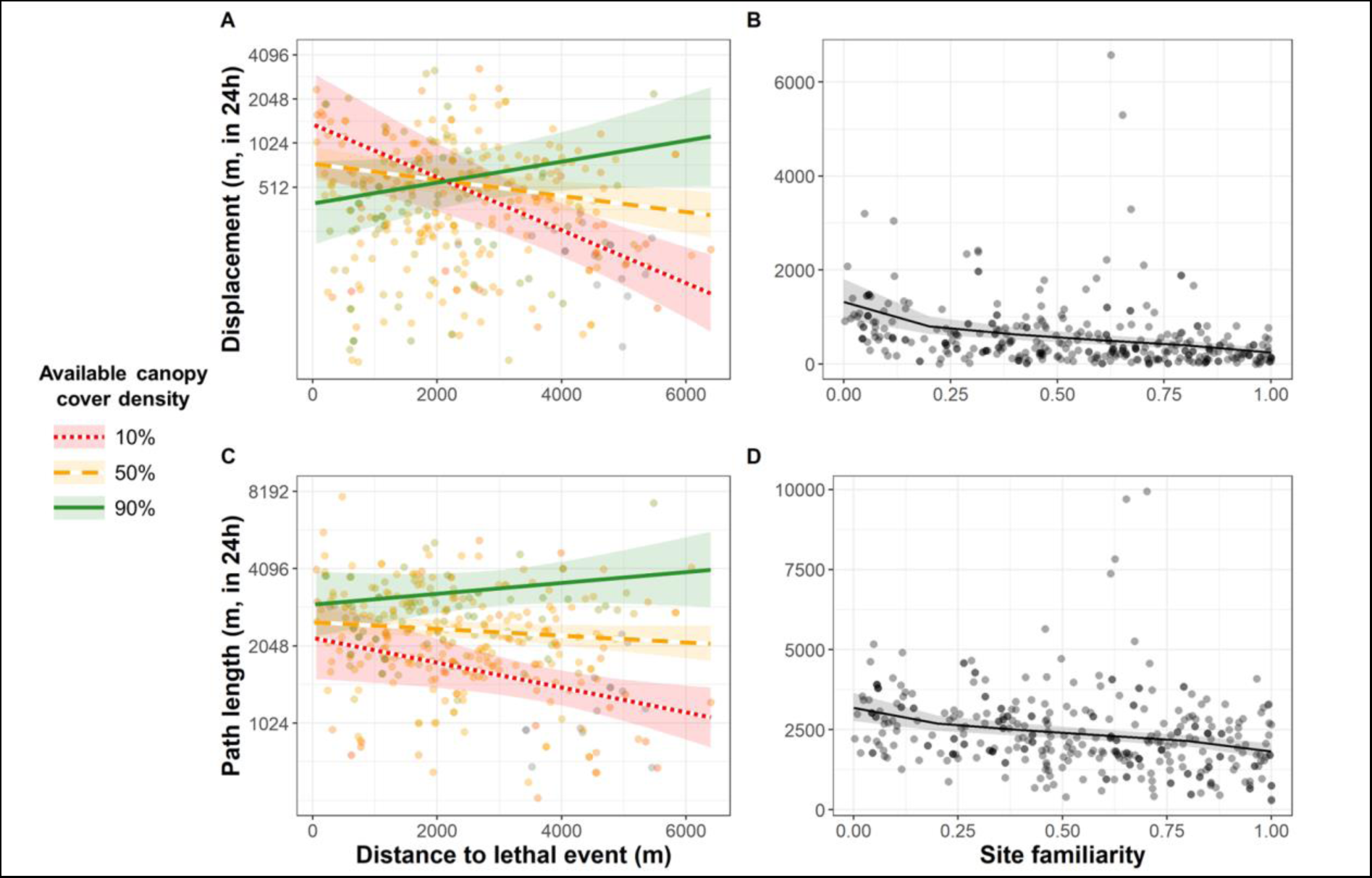
Predictions of red deer movement metrics in the 24 h following a hunting event (**A-B**; displacement, **C-D**; path length). Panels show effects of interaction between distance to the lethal event and available canopy cover density to the red deer at the time of the lethal event (**A-C**), and of site familiarity at the red deer location at the time of the lethal event (**B-D**). Lines with shaded areas represent the model prediction (mean effect with 95% CI), and dots represent observed data. For **A-C**: data are log-transformed for visual purposes; colour/dash patterns refer to three reference canopy cover densities.

### 3.2. Reaction to lethal risk: instantaneous habitat selection response

The model considering two-way interactions between acute spatio-temporal risk and canopy cover density, acute spatio-temporal risk and diel period, and canopy cover density and diel period (i.e. Alternative model 1; Table S8) was retained as the most parsimonious. Red deer avoided acute spatiotemporal risk during the day (Fig.3A, log-RSS [95% CI] = −2.772 [−4.590; −0.955]) and at dusk (Fig.3A, log-RSS [95% CI] = −2.256 [−3.679; −0.834]), while they selected for these areas at night (Fig.3A, log-RSS [95% CI] = 0.926 [0.099; 1.752]). At dawn, movement choices were independent of the acute spatiotemporal risk (Fig.3A, log-RSS [95% CI] = −0.664 [−2.789; 1.460]). In the absence of risk, red deer selected for cover during the day (Fig.3B, log-RSS [95% CI] = 0.517 [0.339; 0.694]) and at dawn (Fig.3B, log-RSS [95% CI] = 2.137 [1.782: 2.492]), and they avoided cover at night (Fig.3B, log-RSS [95% CI] = −1.072 [−1.204; −0.940]) and at dusk (Fig.3B, log-RSS [95% CI] = −2.331 [−2.630; −2.032]). However, selection for cover decreased as acute spatiotemporal risk increased (Fig.3C, slope_log-RSS_ = −1.363). Given the absence of the three-way interaction in the most parsimonious model, this slope of selection for cover as a function of acute spatiotemporal risk was invariable across diel periods.

**Figure 3.**
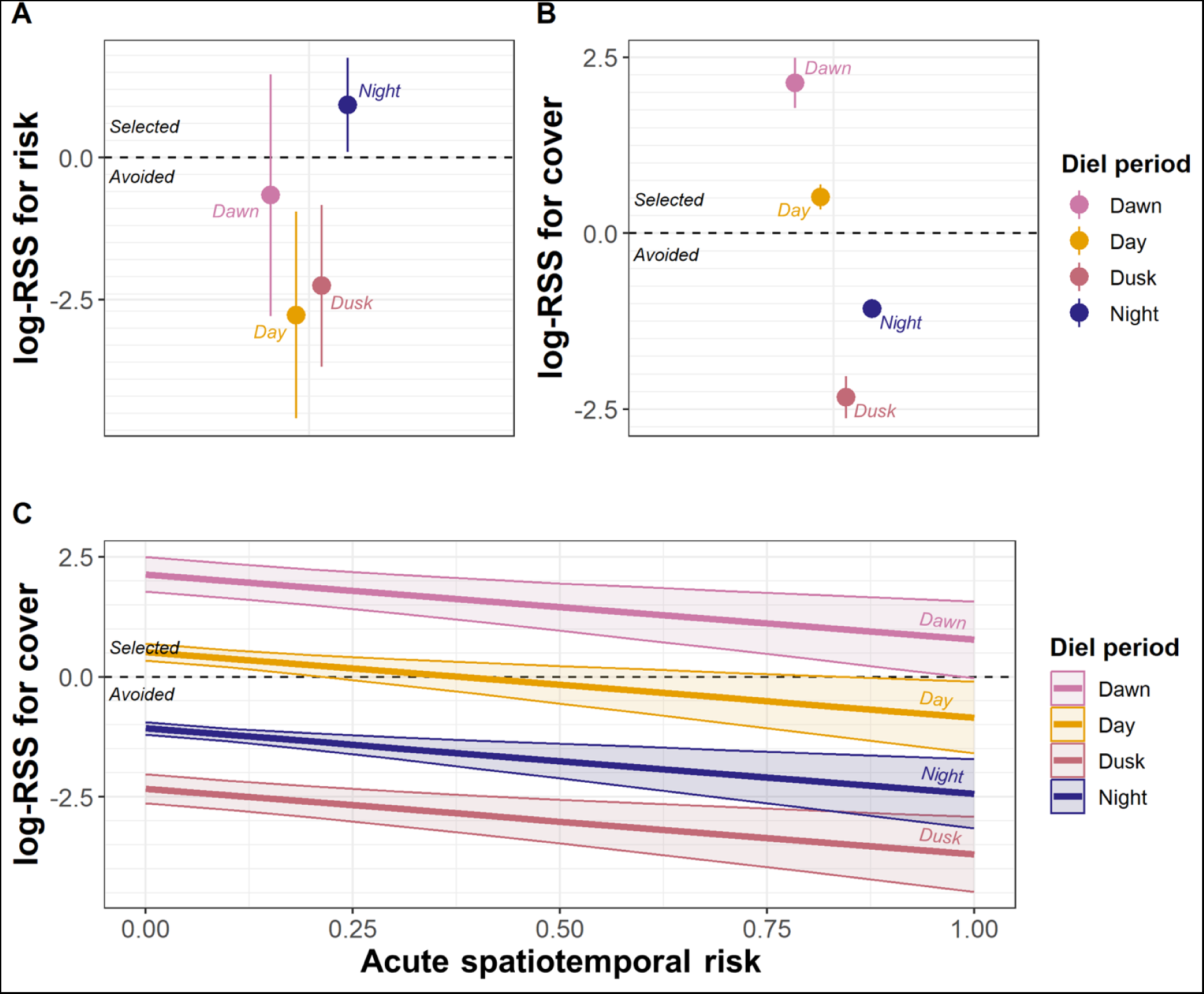
Relative selection strength (log-RSS; Avgar et al. 2017) with 95% confidence intervals for (**A**) acute spatiotemporal risk and for (**B-C**) canopy cover by red deer, at different diel periods (see legend). For **B**, note that the acute risk value was set to 0 in the calculation of log-RSS in order to reflect red deer behaviour in the absence of risk. For **C**, note that diel period affects the absolute level of canopy cover selection (see B), but not its relationship with acute spatiotemporal risk; however, these were plotted for clarity.

### 3.3. Proaction to lethal risk: overall habitat selection behaviour

The model considering the three-way interaction between chronic spatial risk, canopy cover density and diel period, and the two-way interaction between predictable temporal risk and canopy cover (i.e. Alternative model 8; Table S9) was retained as the most parsimonious model. As the level of chronic spatial risk increased – i.e. as the local density of accumulated hunting events throughout the years increases – red deer selected more intensely for cover at dawn (Fig.4A, Table 1), they avoided cover more intensely at night (Fig.4A, Table 1) and even more so at dusk (Fig.4A, Table 1). During the day, red deer selected less for cover as the level of predictable spatial risk increased, but only to a certain level of risk (*circa* 0.75 on a scale from 0 to 1), beyond which cover was used according to availability (Fig.4A, Table 1). In parallel to the annual fluctuation of hunting pressure, red deer also adapted their habitat selection by increasing selection for cover as predictable temporal risk increased (Fig.4B, Δlog-RSS_max-min_ = 0.555). Given the absence of the three-way interaction in the most parsimonious model, this fluctuation throughout the year was invariable across diel periods. Interestingly, selection for cover reached its maximum in mid-September, when the hunting pressure is highest.

**Figure 4.**
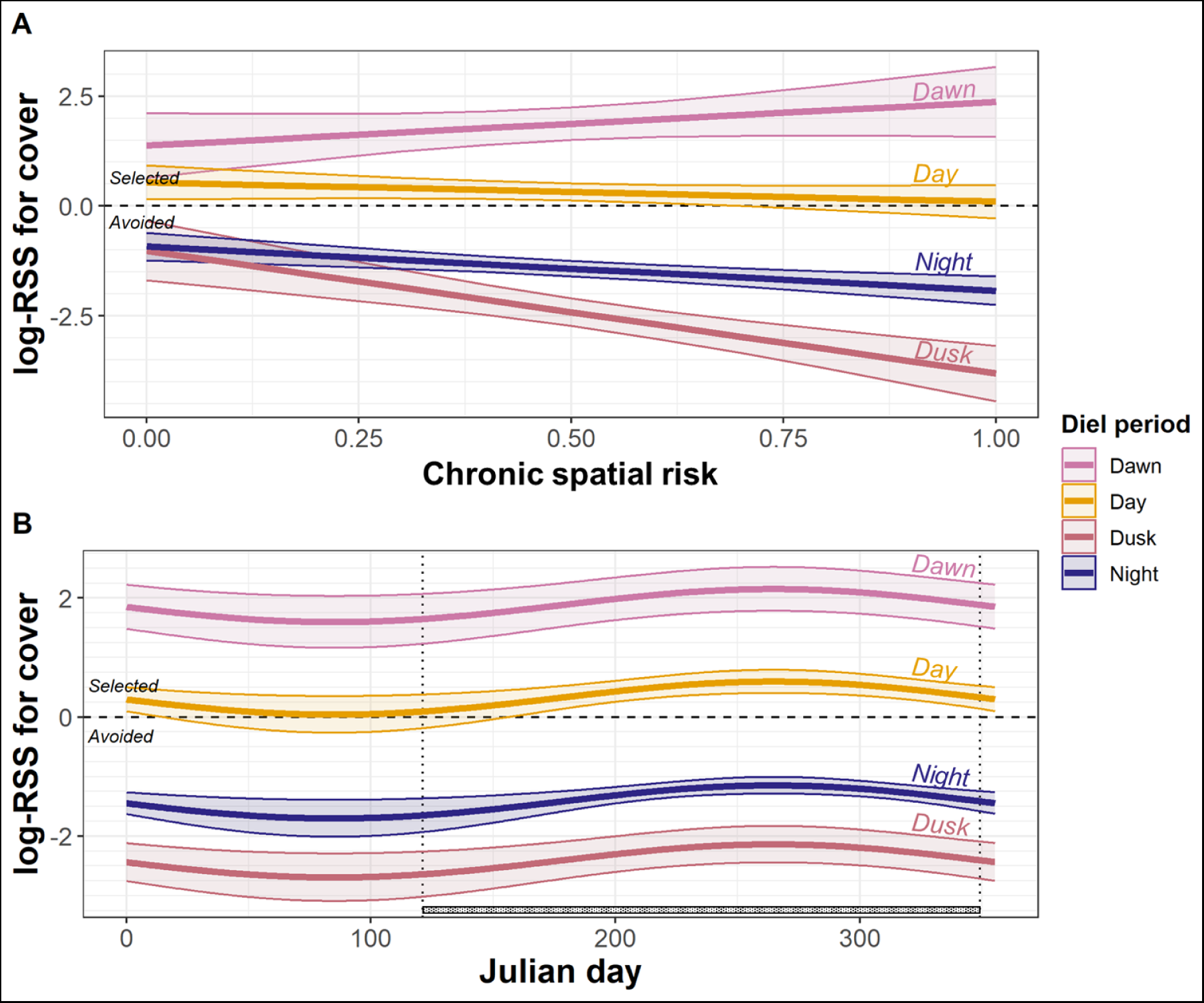
Relative selection strength (log-RSS; Avgar et al. 2017) with 95% confidence intervals for canopy cover by red deer in response to chronic spatial risk (**A**) and to predictable yearly risk (**B**), at different diel periods. For **B**, note that the diel period affects the absolute level of canopy cover selection (see Fig.3B), but not its relationship with predictable yearly risk. The hunting period is indicated with the textured line above the x-axis.

**Table 1.** Relative selection strengths (log-RSS; Avgar et al. 2017) with 95% confidence intervals for canopy cover by red deer in response to low and high chronic spatial risk, for each diel period (dawn, day, dusk, night).

|  | Low chronic spatial risk (set to 0) |  | High chronic spatial risk (set to 1) |  |
| --- | --- | --- | --- | --- |
| Diel period | log-RSS | 95% CI | log-RSS | 95% CI |
| Dawn | 1.378 | [0.641; 2.115] | 2.370 | [1.579; 3.162] |
| Day | 0.540 | [0.154; 0.927] | 0.101 | [-0.280; 0.481] |
| Dusk | -1.017 | [-1.695; -0.339] | -3.812 | [-4.442; -3.182] |
| Night | -0.922 | [-1.239; -0.606] | -1.927 | [-2.248; -1.605] |

## Discussion

Given the pivotal role of ungulates in ecosystems (Pringle et al. 2023, Monk 2024), understanding the landscape-scale dynamics between predators and prey and their potential cascading effects, is key to understand the dynamics and fate of anthropogenic ecosystems (Dorresteijn et al. 2015, Kuijper et al. 2016, Ausilio et al. 2021). Taking advantage of the spatiotemporal scalability of movement paths, and a unique, complete dataset of hunting activities, we investigated antipredator movement behaviour in red deer, in response to human hunting on conspecifics. We aimed to determine how proaction and reaction interact to shape movement behaviours of this prey species at various spatiotemporal scales in their natural environment. By analysing movement behaviour and habitat selection of red deer in response to spatiotemporally heterogeneous predation risk, we showed how canopy and familiarity with the landscape modulate antipredator responses to chronic and acute lethal risk to answer the questions outlined in Fig.1. Specifically, the differential role of canopy cover in proactive and reactive antipredator behaviour revealed an unexpected decisional sequence in red deer: they proactively selected for cover where risk was chronically high (Fig.4A; Q.8) and when risk was predictably high (Fig.4B; Q.9), with adjustments throughout the diel cycle indicating a fine-scale perception of risk. As they selected open high-risk areas at dusk (Fig.2,3; Q.10) – possibly to compensate for lost feeding opportunities throughout the day – they occasionally exposed themselves to acute lethal risk, to which they responded by adjusting movement responses along a freeze-to-escape continuum modulated by the level of risk (distance to the lethal event), surrounding canopy cover and site familiarity (Fig.2; Q.1-4). In particular, deer tended to freeze if they were in covered areas and to escape if they were in open habitats, but less so when the environment was familiar, confirming the importance of memory and cognition in decision-making. As expected, avoidance of acute risk itself was strongest during the day and at dusk (Fig.3A; Q.5,7), when hunting was most probable (88% of hunting events; Fig.S4). However, we did not find selection for cover along the chosen response path (Q.6): the instantaneous habitat selection response showed cover avoidance as acute spatiotemporal risk increased (Fig.3C), indicating that cover at the time of lethal event encounter only conditioned the freeze-to-escape choice (Fig.2A,C).

### Canopy cover and site familiarity modulate the response to acute risk on a freeze-to-escape continuum

Instead of the typical binary dilemma between freezing or escaping in reaction to direct acute risk (Eilam 2005), over 24 h, we found a continuum of behaviours from freezing to escaping, characterised by varying displacements and path lengths, modulated by distance to risk, by the availability of canopy cover, and by site familiarity (Fig.2).

At short distances from risk, cumulated path length remained constant regardless of cover (Fig.2B), confirming a heightened level of alertness after a hunting event as noted by Chassagneux et al. (2019). Under similar conditions, other authors have investigated high reactivity to additional cues (Vanderlocht 2021, Adams et al. 2006); active seeking of information to reassess risk (Blanchard et al. 1991, Hemmi & Pfeil 2010); or an active exploration in search for safer areas (Grignolio et al. 2011). Therefore, red deer response in densely covered habitat seemed to resemble a *restricted movement* response, rather than a strict *freeze* response.

On the other hand, here, exposure to lethal events in habitats with less cover induced more directional movements in deer, as shown in Fig.2A *vs* 2C (red dotted lines), showing similar intercepts but the slope associated with displacement far exceeds the one associated with path length, probably due to an increased perception of risk exposure in this type of environment (Bonnot et al. 2017), or possibly because they can simply be crossed more quickly. Instead, exposure to hunting events in habitats with high cover density induced movements that were always less directional, whether the deer was close to the hunting event or not (Fig.2A *vs* 2C, green solid lines: higher intercept for path length than for displacement, but similar slopes), possibly due to a reduced perception of risk exposure; a need to reassess the risk; or zig-zagging behaviour in order to hinder close pursuits (Caro et al. 2004) or confuse the predator (Humphries & Driver 1967) in habitats with lower visibility and more obstacles. Furthermore, we found a behavioural switch in red deer displacements at *circa* 2 km from lethal events (Fig.2A), suggesting a distance above which red deer decreasingly perceive hazard cues, or respond less as exposure decreased (Ydenberg & Dill 1986). Further research and experimental set-ups are needed to disentangle perception from response (Prugh et al. 2019); nevertheless, the spatial scale of this switching distance was consistent with previously published studies; for example, elk (*Cervus canadensis*) were shown to move from grasslands to forested areas when wolves were detected within 1 km (Creel et al. 2005) and their movement rates decreased as distance to wolves increased up to 5 km, beyond which their movement rates were constant (Proffitt et al. 2009). Finally, both displacement and path length became longer in more densely covered habitats as distance from lethal events increased (Fig.2A,C), indicating in the absence of perceived risk a typical directional use of forested areas and a typical exploitative use of open areas.

In addition, prey species have often been noted to flee risk more often and at greater distances when cover is scarce, when understorey vegetation is scarce, or when far from a refuge (Stankowich and Blumstein 2005). Our measurement of available canopy cover at the time of exposure to lethal events provided a single variable integrating all these various potential elements concurrently used in red deer decision making: the current position in refuge habitat; the nearest distance from refuge habitat; and the availability of refuge habitat. Unlike fleeing, restricted movement behaviour may provide a more adequate response to avoid detection by predators (Endler 1991, De Franceschi et al. 2016), especially in response to visual predators such as humans (Gaynor et al. 2024) in areas with dense canopy cover offering concealment (Smith 1991). We found that strict freezing or hiding, in contrast with restricted movements, may be more likely observed in prey that are physically unable to outrun their predator (e.g. ungulate fawns, Fitzgibbon 1990) or in prey that can effectively be hidden in their environment (e.g. small, solitary, camouflaged and/or woodland species; Smith 1991, Koivula et al. 1995). However, unlike the findings of Chassagneux et al. (2020) and Bojarska et al. (2024) on red deer responses to drive hunts, the restricted movement responses that we observed were not followed by delayed flight within 24 h, and were significantly more likely if the deer were familiar with their surroundings (Fig.2B,D). Restricted movement responses may indeed provide for more safety to sit-and-wait hunting, compared to spatially spread drive hunts (Montgomery et al. 2022). Moreover, the often hypothesised role of familiarity in antipredatory behaviour (Clarke et al. 1993, Forrester et al. 2015) may indeed become more apparent with direct exposure to sit-and-wait hunting events, where risk is acute and precisely localised. Interestingly, high site familiarity reduced displacement (Fig.2B) and path length (Fig.2D) possibly as a consequence of a reduced need for active information seeking (Snyder et al. 1976) and a more effective evasion (Clarke et al. 1993), facilitated by improved navigation abilities in familiar environments (Jang et al. 2019).

### At dusk, red deer are often found in higher-risk environments: compensation for lost opportunities throughout the day or intrinsic clock?

In our study area, dusk appeared to be the most hazardous diel period, as 62% of all red deer hunting events happened in that time (Fig.S4). Interestingly, the riskiest behaviours were also displayed at that time, as red deer avoided cover most strongly at dusk (Fig.3B), and did increasingly so as both acute risk (Fig.3C) and predictable spatial risk increased (Fig.4A). It is important to note that risk taking at dusk was measured with canopy cover avoidance, but the selection for acute risk itself did not increase at dusk (Fig.3A), showing that red deer took risks by selecting open areas, but remained sensitive to acute risk and avoided it whenever hunting was allowed on a daily scale.

Risk taking at dusk may be a compensatory behaviour for lost grazing opportunities after dawn and day largely spent under canopy cover. The effect of accumulated lost feeding opportunities in the previous hours may also explain the difference in risk taking at dawn and dusk: at dawn, red deer have been able to forage abundantly throughout the night, making risk taking less necessary with respect to dusk. Whether this behaviour relates to other physiological or behavioural constraints (e.g. rumen functioning, alternate habitat use in a mixed feeder, intrinsic crepuscularity) requires further investigation. Furthermore, risk taking at dusk could be mitigated or modulated by other factors which were not investigated here (e.g., adjustment of vigilance or grouping behaviour, body condition, reproductive status, weather patterns).

### Diel cyclicity of lethal risk may be the key for red deer to face the food-cover trade-off

The diel cyclicity of risk, as well as the diel biological clock of red deer, appeared to play a crucial role in their antipredator behaviour and their risk taking behaviour. We found here that acute spatiotemporal predation risk was avoided during the day and at dusk, but it was selected at night (Fig.3), indicating a very fine-scale perception of acute risk. Sensitivity to diel cyclicity of risk is rarely considered in studies on reactive antipredator behaviour (but see Valeix et al. 2009, Creel et al. 2014), but is often described in proactive antipredation (Tillmann 2009, Marchand et al. 2015). In our study system, diel predictability of risk modulated the proactive response to chronic spatial risk (‘landscape of fear’; Fig.4A). Specifically, the more risky an area was perceived to be, the more cover was locally selected at dawn, and the more cover was locally avoided at night (Fig.4A), in accordance with the Risk Allocation Hypothesis (Lima & Bednekoff, 1999), predicting high risk taking when hazard is low. However, risk taking at dusk is an exception to this hypothesis, which we speculate might be related to lost feeding opportunities throughout the day (see previous section). In any case, red deer seemed to cope with the spatial trade-off between cover and food, by modulating their spatial behaviour at a fine diel scale. Our results further support the need to integrate schedules of fear (Palmer et al. 2022) to the typically considered landscapes of fear (Gaynor et al. 2019); indeed, as predation risk varies both in time and in space, integrating these axes of variation in analyses rather than investigating them separately may account for much of the observed variability in risk responses and non-consumptive effects (Cusack et al. 2020, Sheriff et al. 2020, Ausilio et al. 2021).

### Proactive choice for cover conditions following reactive behaviour

When acute risk was effectively encountered, red deer used the availability of surrounding canopy cover to evaluate risk and to display a response on a freeze-to-escape continuum (Fig.2A,C), allowing them to quickly avoid the acute risk (Fig.3A), without changes in canopy cover selection along their path (Fig.3C). The proactive and reactive use of cover appear to be different and underpin sequential context-dependent choices, at different spatio-temporal scales. Indeed, although usually described separately, proaction and reaction coexist within the spectrum of behaviours available to prey species and they potentially contribute together to a multistage antipredator sequence (Hemmi & Pfeil 2010), to context-dependent decision-making (e.g. if risk is unpredictable, prey invest only in reaction; Palmer et al. 2022) and even to proactive-reactive trade-offs within a decisional process (see speed-accuracy trade-off; Chittka & Raine 2009, see evidence- *vs* time-based mechanisms; Hawkins & Heathcote 2021) or within behavioural syndromes (see proactive-reactive personality axis: Quinn et al. 2012). Furthermore, even within reactive antipredator behaviours, there are still elements of predictable risk (i.e. diel cyclicity), allowing red deer to modulate their reactions to acute risk through proactive knowledge. In order to keep a clear and ecologically meaningful distinction between proaction and reaction, we recommend accounting for the predictability, and chronicity *vs* acuteness of risk (see Glossary in Palmer et al. 2022), instead of the long *vs* short term nature of the risk (see Broekhuis et al. 2013, Creel et al. 2014) which can be a misleading criterion.

## Conclusions

In this study area and period, it is likely that human hunting was the most important risk for red deer (i.e. humans as a “super predator”: Darimont et al. 2015; approximately 59 human kills per year in the study area *vs* estimated approximately 17 wolf kills per year: Righetti et al. 2019), but the increasing presence of wolves and their different hunting mode (coursing *vs* human sit-and-wait) may also have contributed to shaping red deer space use patterns. The integration of all lethal impacts into future analyses, as well as non-lethal human disturbance, may further contribute to understanding ungulate behaviour in anthropogenic ecosystems.

Our results revealed the ability of red deer to navigate spatiotemporally varying risk in their environment by modulating their movement behaviour at large and fine scales, spatially and temporally, reactively and proactively. Applying a ‘dynamic landscape of fear’ framework (Palmer et al. 2022), we empirically integrated various spatial and temporal scales of risk in our study of antipredator movement behaviour in a hunted ungulate. For the first time in a large herbivore, we described how proaction and reaction fuse in an *antipredation sequence* (Hemmi & Pfeil 2010) of interconnected movement decisions. From these findings, several interesting questions emerge: what are the evolutionary origins of these similarities and differences between proactive and reactive antipredator behaviours? Do proactive behaviours emerge from accumulated reactive behaviours? Or do both behaviours undergo independent natural selection? Further research integrating individual *and* species life-histories may provide novel insights to these intriguing questions.

## Authors’ contributions

**CV:** Conceptualisation; Data curation; Formal analysis; Investigation; Methodology; Software; Validation; Visualisation; Writing – original draft; Writing – review & editing. **BR:** Conceptualisation; Formal analysis; Investigation; Methodology; Software; Validation; Visualisation; Writing – review & editing. **AC:** Formal analysis; Investigation; Software; Visualisation; Writing – review & editing. **SDF:** Data curation; Investigation. **FO:** Data curation; Investigation; Project administration; Resources; Writing – review & editing. **DR:** Data curation; Investigation; Project administration; Resources. **HCH:** Funding acquisition; Project administration; Supervision; Writing – review & editing. **LP:** Funding acquisition; Project administration; Supervision; Writing – review & editing. **FC:** Conceptualisation; Funding acquisition; Project administration; Resources; Supervision; Writing – review & editing.

## Acknowledgements

We thank the South Tyrol Alto Adige Hunters’ Association (Südtiroler Jagdverband/Associazione Cacciatori Alto Adige), especially hunters from the preserves of Lauregno/Laurein and Proves/Proveis; the veterinarians of the Istituto Zooprofilattico Sperimentale delle Venezie (Sezione di Bolzano); and the wildlife technicians of the Wildlife Management Office of the Province of Bolzano for their collaboration. We also thank all the stage students and interns who assisted during captures. We are grateful to the Eurodeer members who provided constructive feedback on a preliminary version of the analysis. This work was financially supported by a Ph.D. scholarship to CV funded by the Centre Agriculture Food Environment - C3A, (University of Trento and the Fondazione Edmund Mach - FEM), and the Stelvio National Park. BR was funded by the Gordon and Betty Moore Foundation (GBMF9881). AC (fully) and FO, HCH, and FC (partially) were funded under the National Recovery and Resilience Plan (NRRP), Mission 4 Component 2 Investment 1.4 - Call for tender No. 3138 of 16 December 2021, rectified by Decree n.3175 of 18 December 2021 of the Italian Ministry of University and Research funded by the European Union – NextGenerationEU (Project code CN_00000033, Concession Decree No. 1034 of 17 June 2022 adopted by the Italian Ministry of University and Research, CUP D43C22001280006, Project title “National Biodiversity Future Center - NBFC”). Views and opinions expressed are, however, those of the author(s) only and do not necessarily reflect those of the European Union or the European Commission, nor can they be held responsible for them.

