## Supplementary information for "Movement responses to lethal risk: an integrative analysis of proactive and reactive antipredator behaviours in a large herbivore"

### Data S1 – Additional information on the study area

Lauregno and Proves have a population density of 23.27 people/km$^2$ and 13.40 people/km$^2$, respectively (ISTAT, 2024). Both municipalities have a well-developed infrastructure network, with numerous forest roads and footpaths across the study area. The climate is alpine, with about 615 mm of mean annual precipitation and mean annual temperatures between -2° and +17°C (1980-2016; https://it.weatherspark.com/). The mammal community includes large herbivores: red deer (*Cervus elaphus*), roe deer (*Capreolus capreolus*), and chamois (*Rupicapra rupicapra*); mesocarnivores: red fox (*Vulpes vulpes*) and badgers (*Meles meles)*; and large carnivores: brown bear (*Ursus arctos*; which was reintroduced in a nearby area between 1999 and 2001, Duprè et al. 1998) and grey wolf (*Canis lupus*; which has been recolonising the area since 2017, Groff et al. 2018).

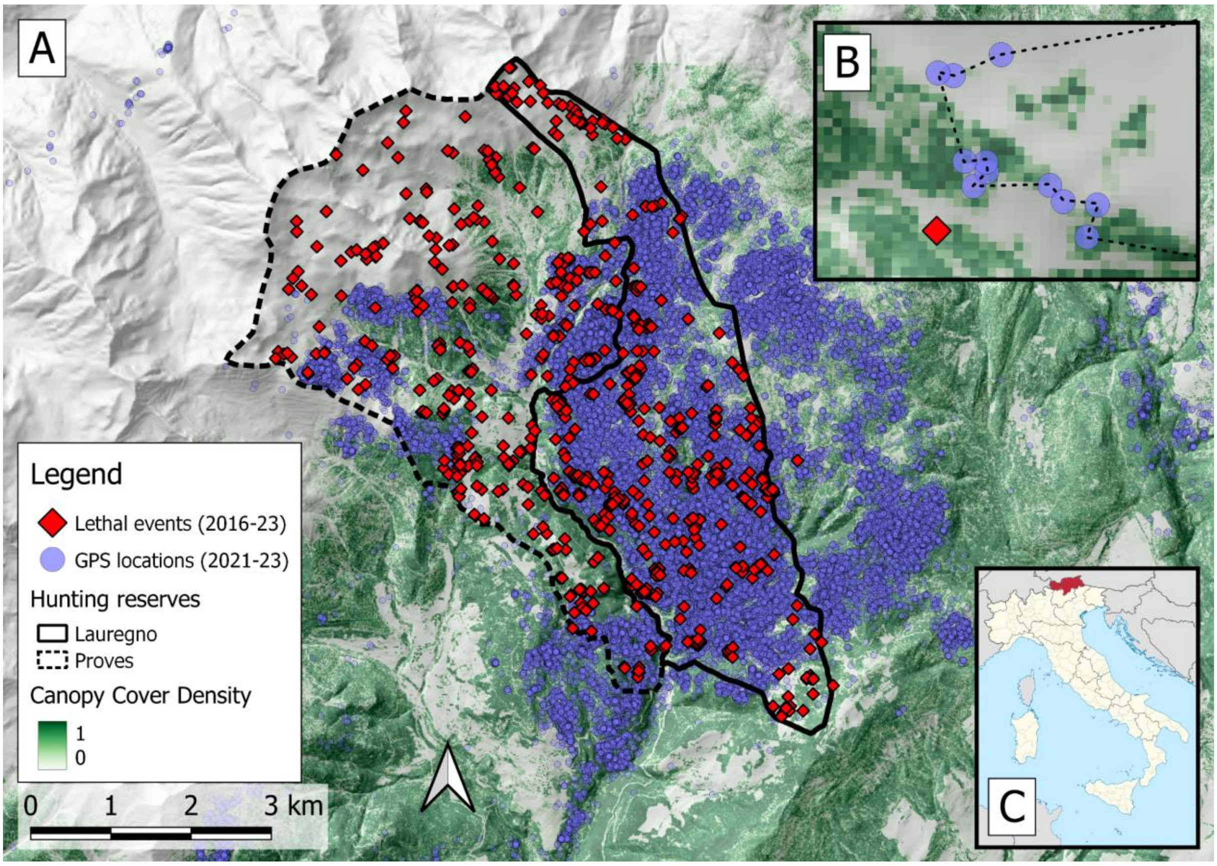


***Figure S1.*** *Plot A: Map of the study area. The lethal events reported by hunters between 2016 and 2023 in the two hunting preserves of Lauregno and Proves (solid and dashed line, respectively) are shown as red diamonds. The GPS locations of red deer monitored within the study area between 2021 and 2023 are shown as purple circles. The LiDAR-derived canopy cover density is shown as a basemap layer. Plot B: An example of red deer movement tracked with a GPS collar in proximity to a reported lethal event. Plot C: Location of the Autonomous Province Bolzano, Italy.*


### Data S2 – List of references for all R packages used


- **adehabitatHR**: Calenge, C. (2006) The package adehabitat for the R software: a tool for the analysis of space and habitat use by animals. Ecological Modelling, 197, 516-519
- **amt**: Signer, J, Fieberg, J., and Avgar, T. (2019). Animal movement tools (amt): R package for managing tracking data and conducting habitat selection analyses. Ecology and Evolution, 9, 880–890.
- **circular**: Agostinelli, C. and Lund, U. (2023). R package 'circular': Circular Statistics. R package version 0.5-0, https://CRAN.R-project.org/package=circular.0
- **DHARMa**: Hartig, F. (2022). DHARMa: Residual Diagnostics for Hierarchical (Multi-Level / Mixed) Regression Models. R package version 0.4.6, https://CRAN.R-project.org/package=DHARMa .
- **glmmTMB**: Brooks, M. E., Kristensen, K., van Benthem, K. J., Magnusson, A., Berg, C. W., Nielsen, A., Skaug, H. J., Maechler, M., and Bolker, B. M. (2017). glmmTMB Balances Speed and Flexibility Among Packages for Zero-inflated Generalized Linear Mixed Modeling. The R Journal, 9(2), 378-400. doi: 10.32614/RJ-2017-066.
- **performance**: Lüdecke, D., Ben-Shachar, M., Patil, I., Waggoner, P., and Makowski, D. (2021). performance: An R Package for Assessment, Comparison and Testing of Statistical Models. *Journal of Open Source Software*, *6*(60), 3139. doi: 10.21105/joss.03139.
- **qdapRegex**: Rinker, T. W. (2017). qdapRegex: Regular Expression Removal, Extraction, and Replacement Tools. 0.7.2. University at Buffalo. Buffalo, New York. http://github.com/trinker/qdapRegex.
- **segclust2d**: Patin, R., Etienne, P. M., Lebarbier, E., Chamaille-Jammes, S., and Benhamou, S. (2020). “Identifying stationary phases in multivariate time series for highlighting behavioural modes and home range settlements.” Journal of Animal Ecology, 89(1), 44-56. doi: 10.1111/1365-2656.13105.
- **spatialEco**: Evans, J. S., and Murphy, M. A. (2023). spatialEco. R package version 2.0-2, https://github.com/jeffreyevans/spatialEco.
- **suncalc**: Thieurmel, B., Elmarhraoui, A. (2022). suncalc: Compute Sun Position, Sunlight Phases, Moon Position and Lunar Phase. R package version 0.5.1, https://CRAN.R-project.org/package=suncalc.
- **terra**: Hijmans, R. (2024). terra: Spatial Data Analysis. R package version 1.7-71, https://CRAN.R-project.org/package=terra.

### Data S3 – Environmental layers

#### Digital elevation model, terrain ruggedness index and slope exposure

We derived the digital elevation model (DEM) for the study area, namely the elevation in metres of the bare ground, from the TINITALY/1.1 dataset (Tarquini et al., 2023), which is available as a 10 m-cell size grid. From the DEM layer we then derived the terrain ruggedness index (TRI), which is defined as the mean altitude difference between a central pixel and all its surrounding cells, and the terrain slope, which is the angle of inclination to the horizontal, using the corresponding functions in QGIS 3.28.6-Firenze (QGIS.org, 2024).

#### Canopy cover layer

We derived the canopy cover from very high-resolution airborne Light Detection and Ranging (LiDAR) data. Survey flights were conducted by plane in 2014, with integration flights in 2018, covering the whole study area. The derived point clouds from the LiDAR scanner were validated, filtered and processed, and a Canopy Height Model (CHM) was provided at a final resolution of 1 m (Autonomous Province of Trento, 2020). From the CHM, we removed all values < 2 m as they represent the shrub layer and therefore not strictly canopy cover. We then derived a canopy cover density by summing all pixels ≥ 2 m and dividing the sum by the unit area (10 m-cell size grid) using the function aggregate from the *terra* R package. The values in the raster layer range from 0 (no cover) to 1 (total canopy cover).

#### Vegetation surveys

Systematic vegetation surveys were conducted as part of a broader project to investigate the emerging ecological functions of large herbivores in the Central Alps in relation to the recent natural recolonization by wolves. We leverage this field sampling information in combination with high-resolution remote sensing data to derive daily food distribution maps. Specifically, vegetation surveys were conducted by randomly placing 1m²-plots throughout a grid system (2 plots per grid cell, over 48 grid cells of 1.5×1.5 km each), covering the hunting reserves of Lauregno-Proveis, and surroundings (Figure S2). In each survey (see field form for vegetation survey; Figure S2), the percentage of area covered by each functional vegetation group (i.e. graminoids, forbs, ferns, shrubs, and mosses/lichens), and their proportions per phenological stage (i.e. newly emergent, budding, flowering, fruiting, mature growth, old growth, cured) were quantified. The average length of plants per functional group and phenological stage, with a specific focus on plants available to ungulates, was thus measured. These vegetation surveys were carried out three times per year during 2022, based on the NDVI greening curves specific for the area (i.e. increasing NDVI: May 1st to July 15th; peak NDVI: July 16th to September 30th; decreasing NDVI: October 1st to October 31st). In 2023, only a subset of vegetation plots were surveyed according to stratified sampling based on the proportion of main available habitat types in the study area (i.e. forest dominated by *Abies alba*, *Larix decidua*, *Picea abies*, *Pinus spp.*, or broadleaf tree species, and grassland < 1400 m, 1400-1800 m, 1800-2100 m, 2100-2400 m, and >2400 m). This was done to ensure that the field sampling proportionally included the major vegetation types according to the local habitat types.

From these vegetation surveys, we calculated the combined biomass (in m³) of newly emergent graminoids, forbs and shrubs (i.e. the fresh green typically consumed by red deer), and we then modelled this food biomass as a function of various remote sensing layers using generalised linear mixed modelling (negative binomial error distribution and a log family, *glmmTMB* R package, *nbinom1* as family). As additive fixed terms, we used NDVI (MODIS 200 m 16 days; Didan, 2021) as a proxy for primary productivity (Pettorelli et al. 2011) yet unable to distinguish between the greenness of ground vegetation (i.e. available forage for red deer) and the greenness of the canopy (i.e. unavailable forage for red deer), in interaction with the cosine-transformed Julian day (peak of cosine at the start of summer), NDVI in interaction with habitat type (as open area or forest with dominance of *Picea abies*, *Abies alba*, etc.), habitat type in interaction with the cosine-transformed Julian day, and of the slope exposure. In order to account for potential discrepancies in biomass estimations between personnel, field operators were added as a random intercept. We then extrapolated this model to all days in our study period for each sampled cell.

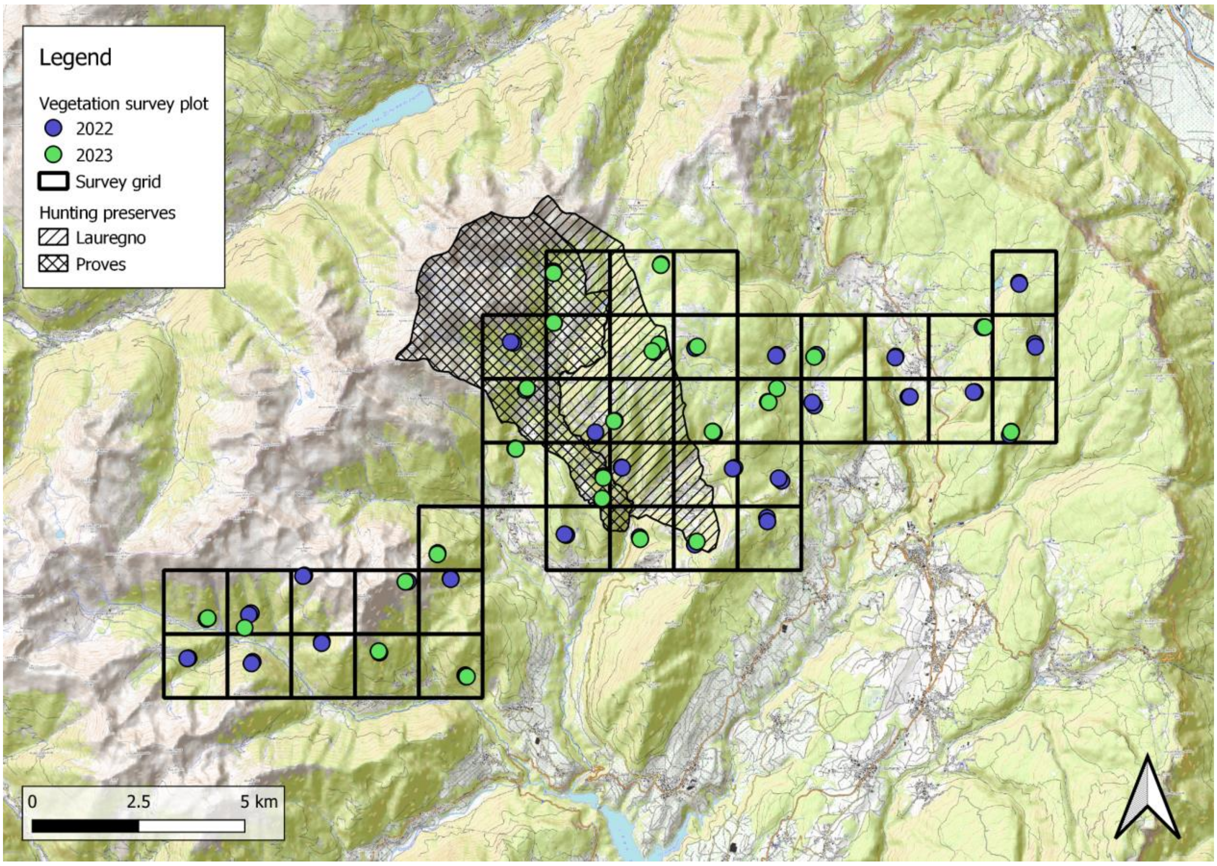


***Figure S3.*** *Surveyed vegetation plot (1 m$^2$, shown as circles) in 2022 and 2023 over the sampling 1.5 x 1.5 km grid. In purple, the plots surveyed in 2022, and in green, the subset of plots surveyed in 2023 according to stratified sampling based on the proportion of main available habitat types. For reference, the two hunting preserves of Lauregno and Proves are shown.*

### Data S4 – Hunting events throughout the the day in Proves and Lauregno

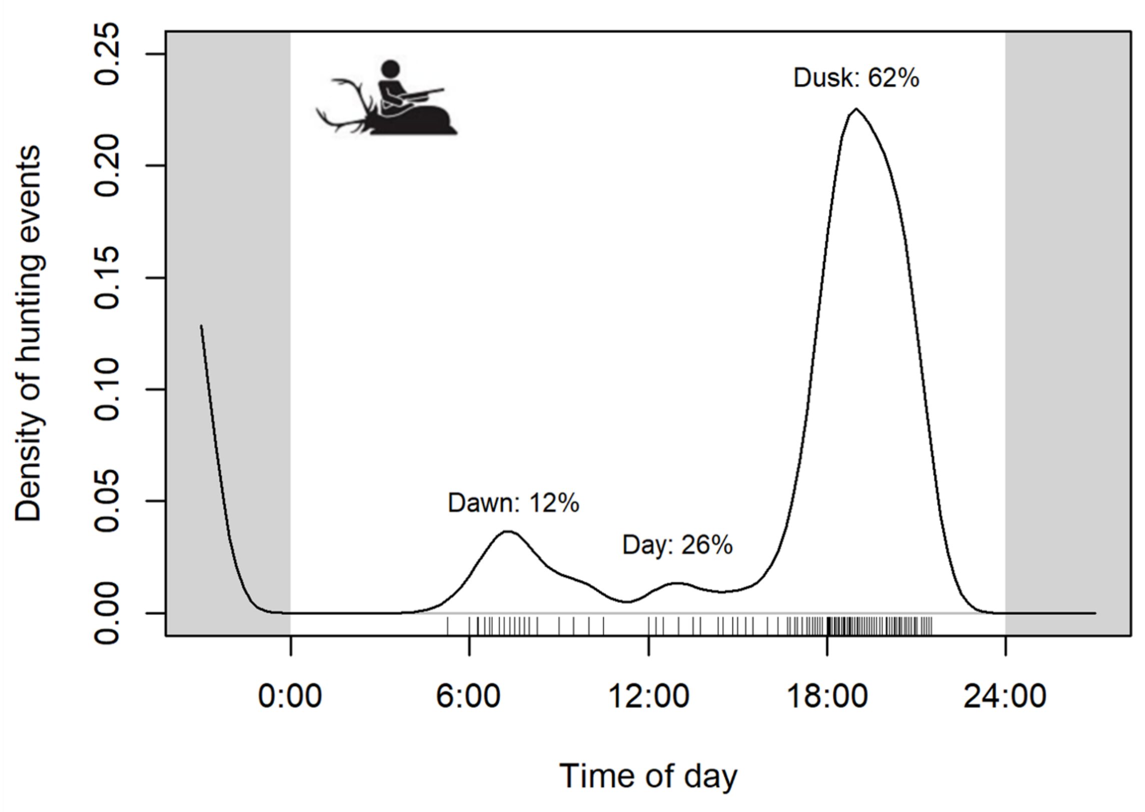


***Figure S4.1.*** *Density of all hunting events throughout the day (time in UCT). We fitted a diel circular kernel density (smoothing parameter of 0.1; circular R package) to the timestamps of all recorded hunting events between January 2016 and November 2023, and we reported the percentage of hunting events happening within each (legal) diel period (Dawn: 12%, Day: 26%, Dusk: 62%).*

### Data S5 – Home range computation

We estimated for each individual the different home ranges as the 95% of the associated movement-based kernel of stationary space use periods. First, we identified phases of space-use stationarity using a bivariate segmentation approach considering the *x* and *y* coordinates of individuals' GPS tracks. We subsampled at 1 fix every 3 h due to computational limits (*segclust2d* R package; minimum length of segment set to 3 days; Patin et al., 2020). For each piecewise stationary period, we then fitted Brownian-bridge based movement kernels (Benhamou & Cornelis 2010; Benhamou 2011) using the *adehabitatHR* R package on the complete GPS track. A brownian bridge was effective only if locations were at no more than 2.5 h from each other. We also filtered out "immobility" considering that movement occurred only if locations were at 50 m from each other. The coefficient of diffusion was estimated based on the movement tracks, and we considered a smoothing parameter of 150 m.

### Data S6 – Output, performance and checks of the displacement model

**Table S6.1.** Model output and performance of the GLMM modelling of red deer displacement as a function of the distance to the lethal event at the time of hunting, interacting with the canopy cover density surrounding the animal (buffer of 500 m) at the time of hunting, and site familiarity, controlling for the local terrain ruggedness at the time and location of hunting, and individuals and years as random effects on the intercept. Test values of single terms involved in significant interactions (-) were not shown due to little to no meaning in our model. Scaled = z-transformation, i.e. scaled to a mean of 0 and a standard deviation of 1.

| **Displacement as a function of …** | | | | |
|---|---|---|---|---|
| *Predictors* | *Estimate* | *SE* | *z* | *p*-value |
| (Intercept) | 6.295 | 0.103 | 60.91 | **< 0.001** |
| Distance to carcass (scaled) | -0.177 | 0.054 | - | - |
| Available canopy cover (500m, scaled after arcsine-square-root transformation) | 0.036 | 0.089 | - | - |
| Familiarity (scaled after arcsine-square-root transformation) | -0.370 | 0.052 | -7.09 | **< 0.001** |
| Local ruggedness (20m, scaled) | -0.029 | 0.057 | -0.50 | 0.618 |
| Distance to carcass x Available canopy cover | 0.178 | 0.052 | 3.41 | **< 0.001** |
| **Random effects** | | | | |
| $\sigma^2$ | 0.58 | | | |
| $\tau_{ind\_id}$ | 0.08 | | | |
| $\tau_{year}$ | 0.00 | | | |
| **Model performance** | | | | |
| Observations | 349 | | | |
| Marginal $R^2$ | 0.345 | | | |
| Conditional $R^2$ | 0.247 | | | |
| AIC | 5055.148 | | | |

**Table S6.2.** Model robustness (outlier test) and normality assumption verification for the "displacement" model (*DHARMa* R package)

| *Statistical test* | *Statistic value* | *p*-value |
|---|---|---|
| Bootstrapped outlier test | Outlier frequency [95% CI]: 0.014 [0.003; 0.026] | 0.640 |
| Asymptotic one-sample Kolmogorov-Smirnov test | Uniformity: 0.043 | 0.535 |

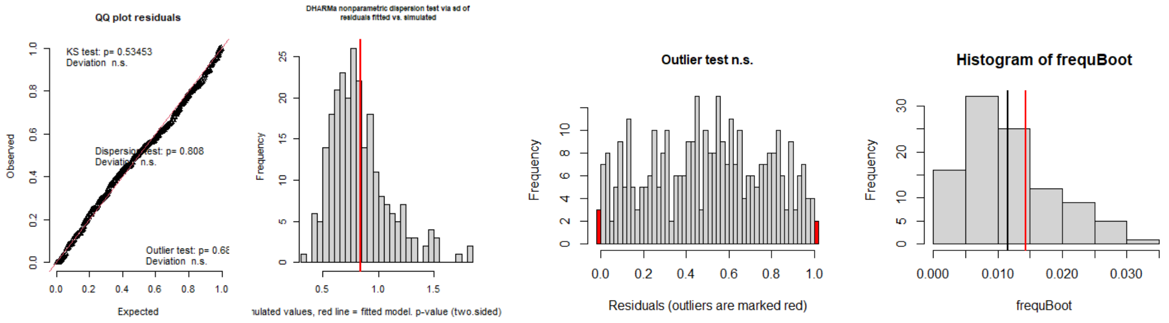


***Figure S6.1.*** *Visual check of the displacement model assumptions (distribution of residuals, QQ-plot of observed vs expected residuals). The plots indicate no violation of the modelling assumptions.*

**Table S6.3.** Collinearity check in the “displacement” model (excluding interactions; *performance* R package)

| *Term* | *VIF* | *VIF 95% CI* | *SE* | *Tolerance* | *Tolerance 95% CI* |
|---|---|---|---|---|---|
| Distance to carcass (scaled) | 1.08 | [1.02; 1.36] | 1.04 | 0.93 | [0.74; 0.98] |
| Available canopy cover (500m, arcsine-square-root transformed) | 1.08 | [1.02; 1.36] | 1.04 | 0.93 | [0.74; 0.98] |
| Familiarity (arcsine-square-root transformed) | 1.06 | [1.01; 1.42] | 1.03 | 0.95 | [0.70; 0.99] |
| Local ruggedness (20m, scaled) | 1.04 | [1.00; 1.54] | 1.02 | 0.96 | [0.65; 1.00] |

### Data S7 – Output, performance and checks of the path length model

**Table S7.1.** Model output and performance of the GLMM modelling red deer path length as a function of the distance to the lethal event at the time of hunting, interacting with the canopy cover density surrounding the animal (buffer of 500 m) at the time of hunting, and site familiarity, controlling for the local terrain ruggedness at the time and location of hunting, and individuals and years as random effects on the intercept. Test values of single terms involved in significant interactions (-) were not shown due to little to no meaning in our model. Scaled = z-transformation, i.e., scaled to a mean of 0 and a standard deviation of 1.

| **Path length as a function of …** | | | | |
|---|---|---|---|---|
| *Predictors* | *Estimate* | *SE* | *z* | *p*-value |
| (Intercept) | 7.766 | 0.038 | 206.73 | **<0.001** |
| Distance to carcass (scaled) | -0.043 | 0.026 | - | - |
| Available canopy cover (500m, scaled after arcsine-square-root transformation) | 0.150 | 0.037 | - | **-** |
| Familiarity (scaled after arcsine-square-root transformation) | -0.122 | 0.025 | -4.95 | **<0.001** |
| Local ruggedness (20m, scaled) | -0.016 | 0.027 | -0.58 | 0.600 |
| Distance to carcass × Available canopy cover | 0.049 | 0.024 | 2.03 | **0.044** |
| **Random effects** | | | | |
| $\sigma^2$ | 0.17 | | | |
| $\tau_{ind\_id}$ | 0.01 | | | |
| $\tau_{year}$ | 0.00 | | | |

| **Model performance** | |
|---|---|
| Observations | 349 |
| Marginal $R^2$ | 0.284 |
| Conditional $R^2$ | 0.268 |
| AIC | 5787.663 |

**Table 7.2.** Model robustness (outlier test) and normality assumption verification for the “path length” model (*DHARMa* R package)

| *Statistical test* | *Test statistic Statistic value* | *p*-value |
|---|---|---|
| Bootstrapped outlier test | Outlier frequency [95% CI]: 0.020 [0.006; 0.023] | 0.360 |
| Asymptotic one-sample Kolmogorov-Smirnov test | Uniformity: 0.061 | 0.145 |

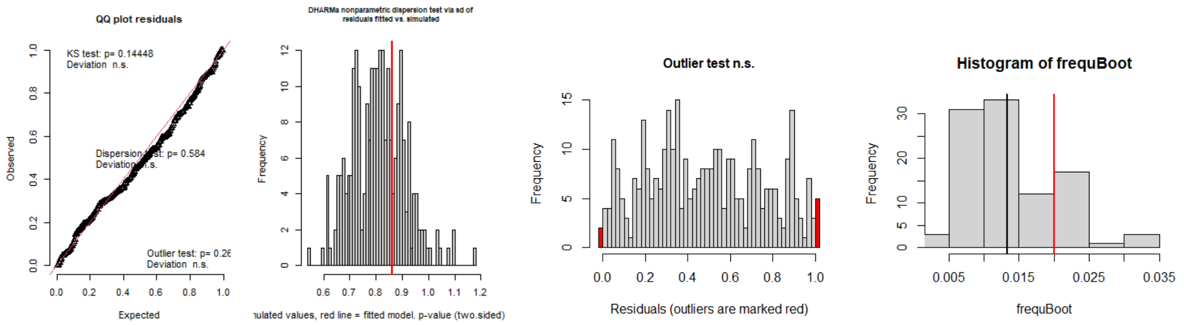


***Figure S7.1.*** *Visual check of the displacement model assumptions (distribution of residuals, QQ-plot of observed vs expected residuals). The plots indicate no violation of the modelling assumptions.*

**Table 7.3.** Collinearity check in the path length model (excluding for interactions; *performance* R package)

| *Term* | *VIF* | *VIF 95% CI* | *SE* | *Tolerance* | *Tolerance 95% CI* |
|---|---|---|---|---|---|
| Distance to carcass (scaled) | 1.12 | [1.04; 1.35] | 1.06 | 0.89 | [0.74; 0.96] |
| Available canopy cover (500m, scaled: arcsine) | 1.11 | [1.04; 1.34] | 1.06 | 0.90 | [0.74; 0.96] |
| Familiarity (scaled: arcsine) | 1.05 | [1.01; 1.48] | 1.02 | 0.95 | [0.68; 0.99] |
| Local ruggedness (20m, scaled) | 1.05 | [1.01; 1.45] | 1.03 | 0.95 | [0.69; 0.99] |

### Data S8 – Model selection for instantaneous habitat selection behaviour

**Table 8.** Comparison of alternative models investigating reactive behaviours to acute lethal risk. AIC = Akaike information criterion, the ΔAIC is obtained by comparing to the most parsimonious model, indicated with an asterisk, and edf = estimated degrees of freedom. In addition to tested variables all models include the control variables as additive terms (+ Food + Altitude + Ruggedness + Step length × (Turning angle + Diel period)) and the individual-step ID as stratum.

| Model | Tested variables | AIC | ∆AIC | edf |
|---|---|---|---|---|
| Null model | / | 79520.10 | 765.95 | 9 |
| Full model | Risk × Cover × Diel period | 78758.50 | 4.35 | 21 |
| Alternative model 1* (simple interactions) | Risk × CoverRisk × Diel period + Cover × Diel period | 78754.15 | 0 | 18 |
| Alternative model 2 (no interactions) | Risk + Cover + Diel period | 79389.60 | 635.45 | 11 |
| Alternative model 3 (no Risk) | Cover × Diel period | 78783.47 | 29.32 | 13 |
| Alternative model 4 (no Cover) | Risk × Diel period | 79503.86 | 749.71 | 13 |
| Alternative model 5 (no Diel period) | Risk × Cover | 79368.52 | 614.37 | 12 |
| Alternative model 6 (no Risk × Cover) | Risk × Diel period + Cover × Diel period | 78765.03 | 10.88 | 17 |
| Alternative model 7 (no Risk × Diel period) | Risk × Cover + Cover × Diel period | 78770.93 | 16.78 | 15 |
| Alternative model 8 (no Cover × Diel period) | Risk × Cover + Risk × Diel period | 79354.02 | 599.87 | 15 |

### Data S9 – Model selection for overall habitat selection behaviour

**Table S9.** Comparison of alternative models investigating proactive behaviours to chronic and predictable lethal risk. AIC = Akaike information criterion, the ∆AIC is obtained by comparing to the most parsimonious model, indicated with an asterisk, and edf = estimated degrees of freedom. In addition to tested variables all models include the control variables as additive terms (+ Food + Altitude + Ruggedness + Step length × (Turning angle + Diel period)) and the individual-step ID as stratum.

| Model | Tested variables | AIC | ∆AIC | edf |
|---|---|---|---|---|
| Null model | / | 79520.10 | 884.25 | 9 |
| Full model | Spatial risk × Cover × Diel period + Temporal risk × Cover × Diel period | 78636.84 | 0.99 | 25 |
| Alternative model 1 (simple interactions) | Spatial risk × Cover + Spatial risk × Diel period + Temporal risk × Cover + Temporal risk × Diel period + Cover × Diel period | 78649.95 | 14.1 | 19 |
| Alternative model 2 (no interactions) | Spatial risk + Temporal risk + Cover + Diel period | 79334.76 | 698.91 | 11 |
| Alternative model 3 (no Temporal risk) | Spatial risk × Cover × Diel period | 78644.42 | 8.57 | 21 |
| Alternative model 4 (no Spatial risk) | Temporal risk × Cover × Diel period | 78769.93 | 134.08 | 17 |
| Alternative model 5 (no Cover) | Spatial risk × Diel period + Temporal risk × Diel period | 79426.13 | 790.28 | 13 |
| Alternative model 6 (no Diel period) | Spatial risk × Cover + Temporal risk × Cover | 79313.17 | 677.32 | 13 |
| Alternative model 7 (no Spatial risk × Cover) | Spatial risk × Diel period + Temporal risk × Cover × Diel period | 78666.30 | 30.45 | 21 |
| Alternative model 8* (no Temporal risk × Diel period) | Spatial risk × Cover × Diel period + Temporal risk × Cover | 78635.85 | 0 | 22 |

| | | | | |
|---|---|---|---|---|
| Alternative model 9 (no Spatial risk × Diel period) | Spatial risk × Cover + Temporal risk × Cover × Diel period | 78686.98 | 51.13 | 19 |
| Alternative model 10 (no Cover × Diel period) | Spatial risk × Cover + Spatial risk × Diel period + Temporal risk × Cover + Temporal risk × Diel period | 79253.80 | 617.95 | 16 |
